# Synergistic mechanistic and deep learning approaches for modeling insect thermal responses and geographic distributions

**DOI:** 10.64898/2026.11.16.766917

**Authors:** Chiara Vanalli, Nina van Tiel, Robin Zbinden, Marie-Moea Geffard Lemaître, Benjamin Kellenberger, Devis Tuia

## Abstract

**Aim:** The Hutchinsonian shortfall, the lack of knowledge about species’ abiotic tolerance, limits our understanding of species responses to environmental change and consequently also the outlining of their distribution in space and time. Such environmental tolerance is commonly inferred with two distinct approaches: *i* ) trait performance studies, which use lab experiments to model species’ functional responses, often overlooking real-world complexity, and *ii* ) species distribution models, which infer environmental suitability from occurrence data, but offer limited causal insights. In this paper, we argue that these two strategies bring complementary information that can be integrated to estimate species–environment relationships more robustly and accurately map species distributions.

**Location:** Global scale

**Time period:** 2000-2019

**Major taxa studied:** Given their exceptional diversity and climatic sensitivity, insects provide a powerful system for studying these responses. We selected key insect species representing *i* ) disease vectors: the yellow fever (*Aedes aegypti* ) and Asian tiger (*Aedes albopictus*) mosquitoes; *ii* ) agricultural pests: the cotton bollworm (*Helicoverpa armigera*) and the fall armyworm (*Spodoptera frugiperda*); and *iii* ) biocontrol agents: the green lacewing (*Chrysoperla carnea*) and the 14-spotted ladybug (*Propylea quatuordecimpunctata*).

**Methods:** We compared predicted insect thermal suitability and distributions derived from two complementary methodological approaches: mechanistic models based on experimentally derived thermal responses of ecological rates, and deep learning-based species distribution models based on occurrence data. We further tested hybrid frameworks integrating mechanistic and correlational information through model ensembling, mechanistic regularization, and mechanistically informed deep learning. We assessed the efficacy of these methods under increasing data scarcity by progressively removing up to 80% of species occurrence records using three strategies: removal from the warmest or coldest parts of species’ thermal ranges, removal from the least-sampled locations based on geographic sampling bias, and random removal as a baseline.

**Results:** Our findings highlight that occurrence-derived thermal suitability closely reproduces the observed distributions of insect. Compared to the mechanistic estimates, these thermal response curves are consistently shifted towards lower temperatures, potentially reflecting niche underfilling. However, when challenged with progressive data removal, correlative methods fail to extrapolate reliably in data-deficient regimes, where mechanistic knowledge becomes increasingly more crucial. Hybrid approaches in turn leveraged complementary information, improving predictive performance and reducing the sensitivity of environmental suitability estimates to sampling biases.

**Main conclusions:** Ultimately, we show how the synergistic integration of experimental ecology with observational biogeography can lead to more accurate and ecologically robust predictions for species distributions and their environmental responses to climatic changes.

## Introduction

Understanding how species respond to environmental conditions remains a focal challenge in ecology [1]. Abiotic factors impact species functional traits [2], population abundances [3], and geographic distributions [4]. These abiotic responses define species invasion potential [5] and their vulnerability and resilience to ongoing climatic change [6], with important implications for ecosystem health and human wellbeing. Yet our knowledge of species’ tolerance to abiotic conditions is still limited, a gap known as the Hutchinsonian shortfall [1]. Ecological data remain incomplete and restricted in spatio-temporal coverage and resolution, often providing only a fragmented representation of species-environment relationships [7]. Consequently, mathematical modeling frameworks that estimate these responses struggle to fully exploit the existing, albeit heterogeneous and biased, knowledge and capture the complex interactions between species and their environment.

The estimation of species’ abiotic responses currently relies on two main approaches: *i.*) trait performance curves from targeted laboratory or field experiments [8, 9], and *ii.*) occurrence-based environmental response curves from species distribution data [10, 11]. These approaches differ fundamentally in the information they exploit and the type of inference they allow. While trait performance curves quantify causal physiological responses and establish mechanistic relations [12, 13], occurrence-based functions statistically infer environmental responses from correlations between abiotic conditions and species presence/absence [11] and do not imply causal links.

The first approach relies on experimental observations to train unimodal (*e.g.*, Brière function [14]) mechanistic performance curves (*MEC*) that increase from a minimum critical limit to an optimum climatic condition, then decline toward the critical climatic maximum. Estimating these cardinal abiotic tolerances allows the identification of operational limits beyond which development, survival, or reproduction is no longer possible [2]. Although these studies provide key insights into the mechanistic abiotic responses of species’ ecological traits, such as growth, survival, development, and fecundity, they are strongly constrained by costly, time-consuming data collection and often consider temperature exclusively [15] while keeping other conditions fixed, therefore limiting their applicability under real environmental conditions.

In contrast, leveraging advances in AI holds the promise of filling biodiversity shortfalls [16]: in our case, deriving response curves from species occurrences has greatly benefited from recent advances in data collection and species identification technologies, using images, videos, and sound recordings, as well as from the growing popularity of species occurrence databases (*e.g.*, 3.6 billion species records in the Global Biodiversity Information Facility [GBIF]) [17, 18]. Modern correlative Species Distribution Models (SDMs) increasingly leverage these extensive data sources to identify statistical patterns between environmental conditions and species occurrences to estimate species environmental suitability [19, 20]. In particular, deep learning (*DL*)-based SDMs have recently shown great potential to integrate heterogeneous and vast amounts of data and extend capabilities of correlative studies [21]. The environment-suitability functions inferred by (Deep)SDMs provide information regarding critical abiotic limits. but can suffer from incomplete sampling of the environmental gradient, a problem known as niche truncation [22]. These limitations are exacerbated by spatial and temporal gaps in species occurrence data, meaning that even common and well-sampled species may remain incompletely characterized across their full distributional range. Moreover, species occurrence datasets suffer from multiple biases: for instance, occurrence data are usually more abundant in higher-income and more accessible regions, which rarely correspond to biodiversity hotspots with high monitoring priority [23, 24].

Hybrid approaches combine mechanistic knowledge and data-informed correlative understanding [25, 26]. They have successfully been shown to overcome single-approach limitations in many fields, including climatic sciences [27, 28] and biomedical research [29]. Hybrid approaches using deep learning in particular have received increased traction, thanks to their flexibility and ability to join correlative power and process understanding, leading to the development of process-informed neural networks [30, 25]. These methods exploit the current understanding of system dynamics, often achieved *in silico*, in combination with correlative information carried by real-world observations. Therefore, the benefits of hybrid methods are twofold: on the one hand, they ground correlative approaches to process-based system knowledge, restricting their predictions to a sensible output space; on the other, they expand what is missing (*i.e.*, not modeled by procedural rules) in current mechanistic understanding of the system. For instance, assumptions underlying process-based models developed *in silico* may not hold under real-world conditions, potentially leading to biased estimates if these additional factors are not adequately accounted for. Despite their potential, hybrid approaches remain underexplored in ecology, especially to study abiotic responses and integrate trait-based ecological theory with species occurrence records.

Among all taxa, insects provide one of the best-studied systems for understanding species’ abiotic responses. Their responses have been extensively characterized through both experimentally derived climate performance curves for key ecological traits [31, 13] and environmental responses inferred from species occurrence observations (*e.g.*, more than 320 million insect records are available from GBIF only). As the most species-rich eukaryotic class and highly dependent on environmental conditions due to their ectothermic nature, insects are particularly sensitive to climate change [32]. Therefore, they are often chosen as sentinel species to provide broader insights into biodiversity response to global change [33]. Importantly, insects provide essential ecosystem services, such as biological control, nutrient cycling, and pollination [34], but also pose significant threats to global health and food security by serving as vectors of infectious diseases [35] and causing substantial damage to agricultural crops [36]. The mechanistic and correlative approaches described above have been directly compared to study the thermal biology of *Aedes*, *Anopheles*, and *Culex* mosquito species, with a strong level of agreement regarding the identification of thermal minima and optima, but not of their thermal maxima, which could not be determined using SDM-inferred thermal curves due to extrapolation issues outside the thermal region where mosquito observations were available [37].

In this work, we further exploit the relative strengths of these complimentary approached and address two fundamental research questions that are addressed in this work: How can ecological knowledge be effectively integrated with large-scale observational datasets to accurately estimate insect response to environmental conditions and map their geographic distributions? What are the respective opportunities brought by the mechanistic and correlative approaches to advance species distribution modeling?

In this work, we address these questions by selecting key insect species, which are disease vectors (yellow fever, *Aedes aegypti*, and asian tiger, *Aedes albopictus*, mosquitoes [38]), major agricultural pests (cotton bollworm, *Helicoverpa armigera* [39], and fall armyworm, *Spodoptera frugiperda* [40]), and effective biocontrol agents (common green lacewing, *Chrysoperla carnea* [41], and fourteen-spotted lady beetle *Propylea quatuordecimpunctata* [42]). These species come with both well-characterized ecological trait thermal response curves and abundant occurrence records. We compiled literature knowledge regarding the thermal response of species ecological rates (i.e., fecundity, immature survival, development, and adult lifespan) and estimating a process-based species suitability. We then examined data-driven species thermal response, based on species presence records through a deep learning-based correlative approach. Last, we proposed and tested multiple hybrid frameworks to integrate the mechanistic and data-driven approaches together (model ensembling, mechanistic regularization, and mechanistically informed deep learning, see Figure 1).

**Figure 1:**
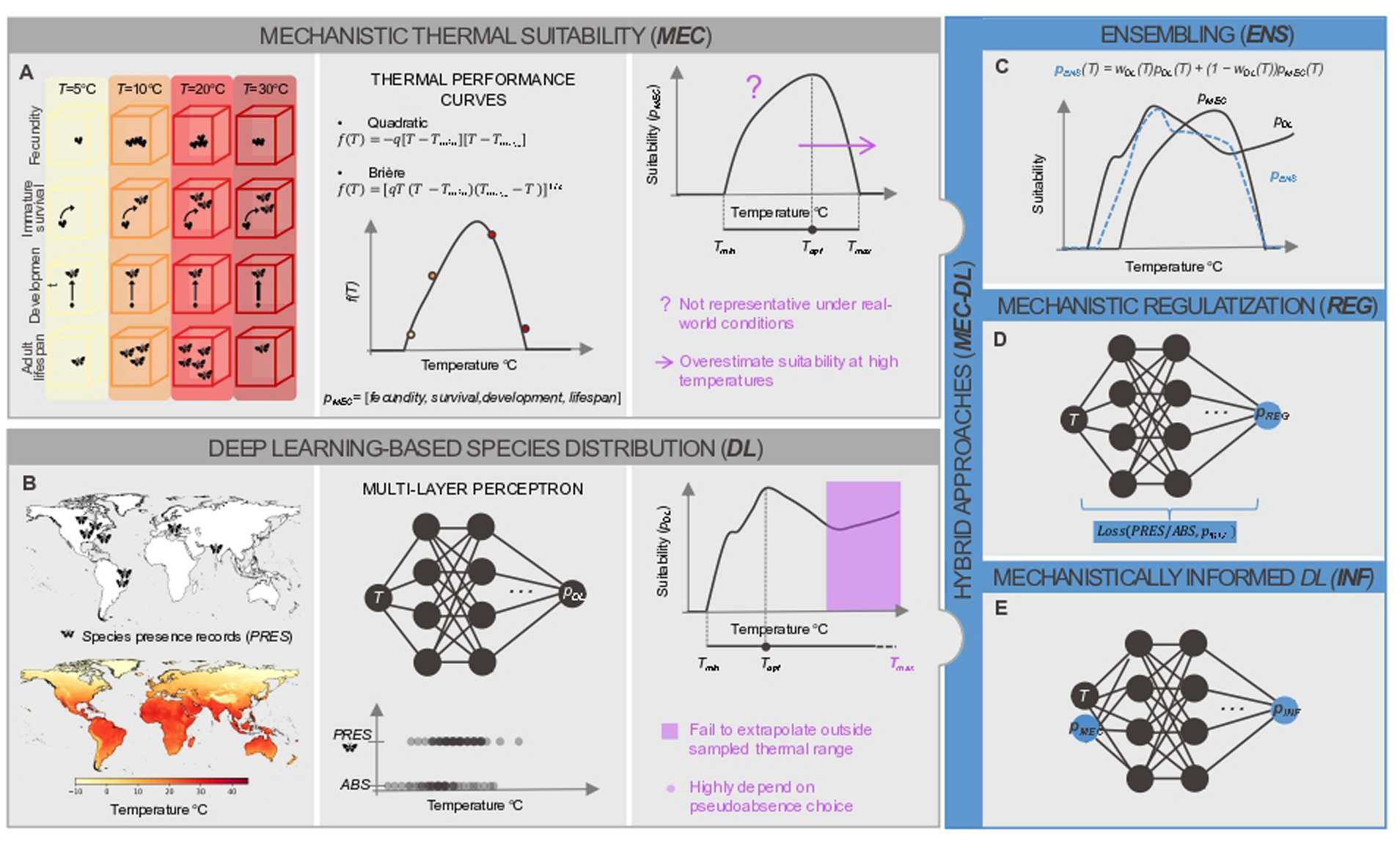
Schematic representation of (A) mechanistic and (B) deep learning-based suitability models (in grey), and of various hybrid approaches: (C) ensembling, (D) mechanistic regularization and (E) mechanistically informed deep learning (in blue). The different input data, modeling scheme, and outputs are described for the mechanistic and deep-learning approaches, with the respective drawbacks highlighted in purple. For further details, please refer to the Materials and Methods section.

Despite the longstanding call from the scientific community to develop correlative-mechanistic integrated frameworks [43, 26], such approaches remain limited. Our study helps fill this gap by showing how integrating experimental ecology with observational biogeography can improve the accuracy and ecological relevance of predictions of species distributions and environmental tolerances across a wide range of insect species. Importantly, this work bridges different disciplines (empirical ecology and deep learning), data sources (experiments and species records), and modeling techniques (climate performance curves and species distribution models) to generate a more robust and unified understanding of insect thermal biology.

## Material and Methods

### Data

#### Insect species

We consider six insect species as case study, selected based on their broad relevance across different ecological fields, the availability of experimental observations regarding their ecological traits from the literature, and the number of available occurrence records from GBIF. We selected mosquito species *Aedes aegypti* and *albopictus* [38], which are important vectors of human infections, moths species *Helicoverpa armigera* [39] and *Spodoptera frugiperda* [40], which damage agricultural crop production, and the green lacewing *Chrysoperla carnea* [41] and ladybug *Propylea quatuordecimpunctata* [42], which are effective biological control agents of aphid pests.

#### Experimental observations of insect thermal biology

Mechanistic thermal suitability requires thermal performance information of the different ecological traits of a species, which we compiled from available knowledge from the literature. We gathered data regarding the thermal dependence of immature survival (%), fecundity (*eggs/*(*female day*) or *eggs/*(*female cycle*)), development rate (1*/day*), and adult lifespan (*day*) tested in the lab under controlled conditions (Table A1) [44, 12, 45, 46, 47, 48, 49]. These data are obtained through experiments performed at constant temperature conditions in incubation chambers using laboratory insect colonies. The gathered data, or the respective parametrized thermal performance curves, are used to establish the mechanistic thermal response of the insect species.

#### Species occurrence records and pseudoabsences

Correlative approaches like the SDMs considered below need observation-specific records of abiotic conditions. We first obtained presence-only data from GBIF for the selected insect species with georeferenced coordinates available in the time period from 2000 to 2019 [50, 51, 52, 53, 54, 55]. Given the importance of seasonality for insects, we accounted for the month of observation by pairing each insect occurrence record with the average monthly temperature (*^◦^C*), calculated over the 2000-2019 period, globally available at 1 *km*^2^ of resolution from CHELSA [56]. The use of monthly global temperatures allows for the estimation of insect suitability for each month of the year providing key insights into intra-annual suitability fluctuations. After download, occurrences went through a series of filtering steps:

- We retained occurrences with a coordinate uncertainty lower than 1 *km*, and kept complete temporal information (month and year).
- We filtered for species-specific activity season, *i.e.*, we excluded species records that belong to dormancy (Figure A1 and Table A2) for *Ae. albopictus*, *H. armigera*, *C. carnea*, and *P. quatuordecimpunctata*, which are species with literature evidence of undergoing a dormant state.
- Species records belonging to the activity season were then approximated using species-specific photoperiod requirements, below which dormancy is initiated, specifically 9 hours of daylight for *Ae. albopictus* [57], 12 hours for *H. armigera* [58], 13 hours for *C. carnea* [59], and 14 hours for *P. quatuordecimpunctata* [60]. Daily photoperiod was estimated using the Python package Astral (version 3.2), based on the calculated times of sunrise and sunset for each location and observation time.

While the datasets describe where each species has been found, they lack records of species absences, which are required for SDMs to discriminate suitable from unsuitable environmental conditions. Common approaches rely on generating of artificial absence data, usually called pseudoabsences or background points depending on the methodology [61, 62, 63]. We generated 10,000 pseudoabsences for each species, randomly sampling from other insect presence records reported in GBIF [64] (i.e., target-group background points) and stratifying them across a thermal range from -25 to 45 *^◦^C*. For target-group background points, the geographic bias in species presence records is reflected in the selection of pseudoabsences. Instead, the environmentally stratified background points take into account the full thermal gradient and the global scope of our study. In both cases, we considered only pseudo-absences at least 1 *km* distant from presence records of the target species.

### Models

#### Mechanistic thermal suitability (*MEC* )

Based on the experimental observations of insect thermal biology (Table A1), thermal performance functions for each investigated ecological trait (immature survival, fecundity, development rate and adult lifespan) can be established (Figures 1A and A3). These functions are typically unimodal [65], with performance increasing from a minimum threshold (*T_min_*) to an optimal condition (*T_opt_*) and then declining rapidly with a null performance beyond a maximum extreme condition (*T_max_*). Depending on the analyzed ecological trait, different temperature-dependent (*T* ) mathematical functions are commonly used, such as a symmetric quadratic curve –*q*[*T* –*T_min_*][*T* –*T_max_*] [14] or the right-skewed Brière function [*qT* (*T −T_min_*)(*T_max_−T* )]^1^*^/^*^2^ [14], with *q* being a scaling constant. Other functional forms can also well describe the thermal response of specific traits, such as the Logan model [66] and the extreme-value function [67]. When the curves were already parametrized in published studies, we directly implemented the available thermal functions; otherwise, we trained the described functional forms (parameters *q*, *T_min_*, and *T_max_*), with a sum of squared residuals loss function (Table A1).

To integrate the different mechanistic trait models into a single value of occurrence probability, we first normalized each trait response function to a maximum value of 1. We then averaged the normalized trait functions and obtained a suitability index ranging from 0 to 1. This procedure was adopted instead of relying on classical population abundance [68] because species environmental suitability is largely unrelated to abundance [69]. A region may provide suitable conditions only for a particular ecological process of the entire insect life cycle, resulting in positive suitability, while simulated population abundance remains null due to spatial independence assumptions.

#### Thermal suitability from deep learning-based species distribution modeling (*DL*)

As an SDM, we implement a Multi-Layer Perceptron (MLP), an artificial neural network primarily composed of fully-connected layers and non-linear activation functions (Figure 1B). MLPs have been widely applied in species distribution modeling due to their ability to capture complex, non-linear correlational relationships between species distributions and environmental predictors, and are well-suited for processing tabular data [70, 71, 72]. We implemented the same MLP architecture across different species, accepting a one-dimensional input for the temperature and predicting species suitability using two hidden layers of 50 neurons each and one output layer. As nonlinearities, ReLU activation functions are used and a final sigmoidal transformation is employed to predict probability values between 0 and 1. Once trained, the MLP can be used to predict spatially-explicit species suitability by performing prediction at all locations on the spatial grid; crucially, it can further be employed to infer how suitability changes along thermal gradients to estimate the species’ thermal response. MLPs were trained using the species presence records and pseudoabsences described above, using a binary cross-entropy loss and Adam optimizer [73] with a learning rate of 0.001 and a maximum of 100 epochs for training. Early stopping was applied based on the cross-entropy loss evaluated on the validation set (see model performance evaluation, comparison, and sensitivity section below).

### Hybrid approaches (*MEC-DL*)

With hybridization approaches (*MEC-DL*) we expect to bridge the complementary strengths of *MEC* and *DL*, by synergizing process-based ecological knowledge with occurrence-driven patterns (Figure 1C-E). We present three model variants combining mechanistic and correlative methodologies, which differ depending on the intended objective and emphasis on either component.

- **Model ensembling (*ENS_MEC−DL_*)**. The outputs of process-based thermal suitability, *p_MEC_*, and occurrence-driven MLP, *p_DL_*, can be seen as complementary descriptions of the same underlying process, akin to specialized experts based on distinct sources of information (Figure 1C). Consequently, combining their predictions within a mixture-of-experts framework may provide a more accurate estimate of species suitability [74]. Because the occurrence-driven *DL* is trained on opportunistically collected observations, its predictions are expected to be more reliable within the range of thermal conditions represented in the training data. In contrast, *MEC* remains applicable across the full thermal gradient, providing a robust basis for extrapolation beyond the observed conditions. Despite a broader thermal coverage, it is worth highlighting that experimental observations often lack measurements at the lowest temperatures.(Figure A3). In *ENS_MEC−DL_*, we combined the two suitability estimates using a temperature-dependent ensemble weighting scheme: *p_ENS_*(*T* ) = *w_DL_*(*T* )*p_DL_*(*T* ) + (1 *− w_DL_*(*T* ))*p_MEC_*(*T* ), where *w_DL_*(*T* ) represents the confidence assigned to the occurrence-derived suitability at temperature (*T* ) estimated from the local density of training presence records along the thermal gradient (using a moving window of 5*^◦^C* and a step of 0.1*^◦^C*). The resulting ensembled suitability depends more strongly on *p_DL_* in regions of the thermal space that are well-represented by the training data, while progressively relying on *p_MEC_* in regions where observations are sparser or entirely absent.
- **Mechanistic regularization (*REG_MEC−DL_*)**. In this variant, the *DL* model is trained with a dual objective: besides the observation status (presence or pseudo-absence), the model is further constrained to minimize the distance between its prediction and the one of the *MEC* model (Figure 1D). To this end, the mechanistic information remains fixed and is used within an additional loss term: *L_REG_* = *L*(*p, p_DL_*) + *λL*(*p_MEC_, p_DL_*), where *L* represents the binary cross-entropy loss and parameter *λ ∈* (0, 1] determines the strength of the regularization [25]. For our main model comparison, we fixed *λ* at 0.3 and investigated its effects on model performance through a species-specific sensitivity analysis (Table A3).
- **Mechanistically informed deep learning (*INF_MEC−DL_*)**. A third approach casts mechanistic ecological knowledge as an additional input to the DL method (Figure 1E). The inclusion of this extra mechanistic input allows the correlative model to incorporate process-based information into the learning process. To do so, *p_MEC_*is first computed withe mechanistic approach and then included as an additional input feature to the MLP alongside the temperature (the MLP’s first layer contains an additional set of weights to accommodate this extra input).

### Model evaluation, comparison, and sensitivity

#### Data splits

While *MEC* models are trained on experimental observations from the literature and only tested on the species occurrence records, *DL* and *MEC − DL* are entirely trained, validated, and tested on the latter. Species presence records and pseudoabsences are split into 80% train, 10% validation, and 10% test sets using spatial blocks of five degrees latitude/longitude. This data split reduces the risk of overestimating model performance due to spatial autocorrelation and provides a more realistic assessment of model transferability to low-sampled regions. Importantly, the 10% test set was used for the comparison of model performance across different approaches and hybridization techniques. To quantify the effect of sampling uncertainty on model performance and predicted suitability outputs, the train-validation-test partitioning procedure was repeated 100 times through random resampling.

#### Model performance

We assessed the performance of the different models on the test set by computing the Area Under the Receiver Operating Curve (*AUC_ROC_*), which summarizes the trade-off between sensitivity and specificity across all possible suitability thresholds. To binarize suitability outputs for the computation of the latter metric, we used the optimal threshold that maximizes the true skill statistic (*TSS*), a threshold-dependent metric, for the validation dataset.

#### Model robustness to degraded species occurrence records

We performed a data degradation analysis to assess whether data scarcity of species occurrence records might affect the difference in model performance and predicted species suitability between hybrid approaches and *DL* alone. For each species, we progressively removed up to 80% of presence records according to three degradation strategies: *i.*) along the thermal gradient by removing occurrences from either the warmest (upper thermal limit) or the coldest (lower thermal limit) parts of the species distribution; *ii.*) according to the geographic sampling bias estimated from all insect records [64] (Figure A2), by removing occurrences from the least-sampled locations; and *iii.*) randomly as a comparison baseline. The first two degradation strategies better reflect the effects of sampling limitations and range truncation commonly encountered in biodiversity datasets and represent realistic scenarios where the integration of mechanistic ecological knowledge can play a crucial role. Note that pseudoabsences remain unaltered in this analysis. As described above, we performed 100 resampling iterations of the train–validation-test blocks and retrained each model for every degradation technique and removal step. While model performance was evaluated across the full gradient of data completeness, we then selected 75% of data completeness to compare the thermal suitability outputs from the different modeling approaches.

#### Model simulation and output analysis

We compared the inferred thermal responses across the multiple tested modeling approaches and calculated the respective optimal temperatures and operational thermal ranges. Thermal optima (*T_opt_*) were defined as the temperature at highest species suitability (*argmax*(*p*)), while suitable thermal ranges were identified as temperatures at which suitability is greater than a minimum threshold, fixed at 10*^−^*^4^. Additionally, using global temperature records (see Species occurrence records and pseudoabsences section), we mapped global species suitability from the modeling approaches and computed the respective differences, highlighting geographic regions of agreement and disagreement across the multiple models.

## Results

### Predicting insect thermal suitability and distribution

For the mechanistic approach, we brought together available knowledge from published literature studies on species’ ecological trait thermal responses and parametrized the thermal functions of the traits for which only experimental observations were available (Figure A3). Aggregating then the thermal response of fecundity, immature survival, development, and adult lifespan, we obtained the suitability for each analyzed insect species along the thermal gradient (Figure 2). These functions are consistently unimodal across species and left-skewed towards higher temperatures. Thermal optima that correspond to highest species suitability are in the range of [25.0, 28.1] *^◦^*C, specifically 28.1*^◦^*C for *Ae. aegypti*, 25.6*^◦^*C for *Ae. albopictus*, 26.0*^◦^*C for *H. armigera*, 25.0*^◦^*C for *S. frugiperda*, 25.8*^◦^*C for *C. carnea*, and 25.7*^◦^*C for *P. quatuordecimpunctata*, respectively.

**Figure 2:**
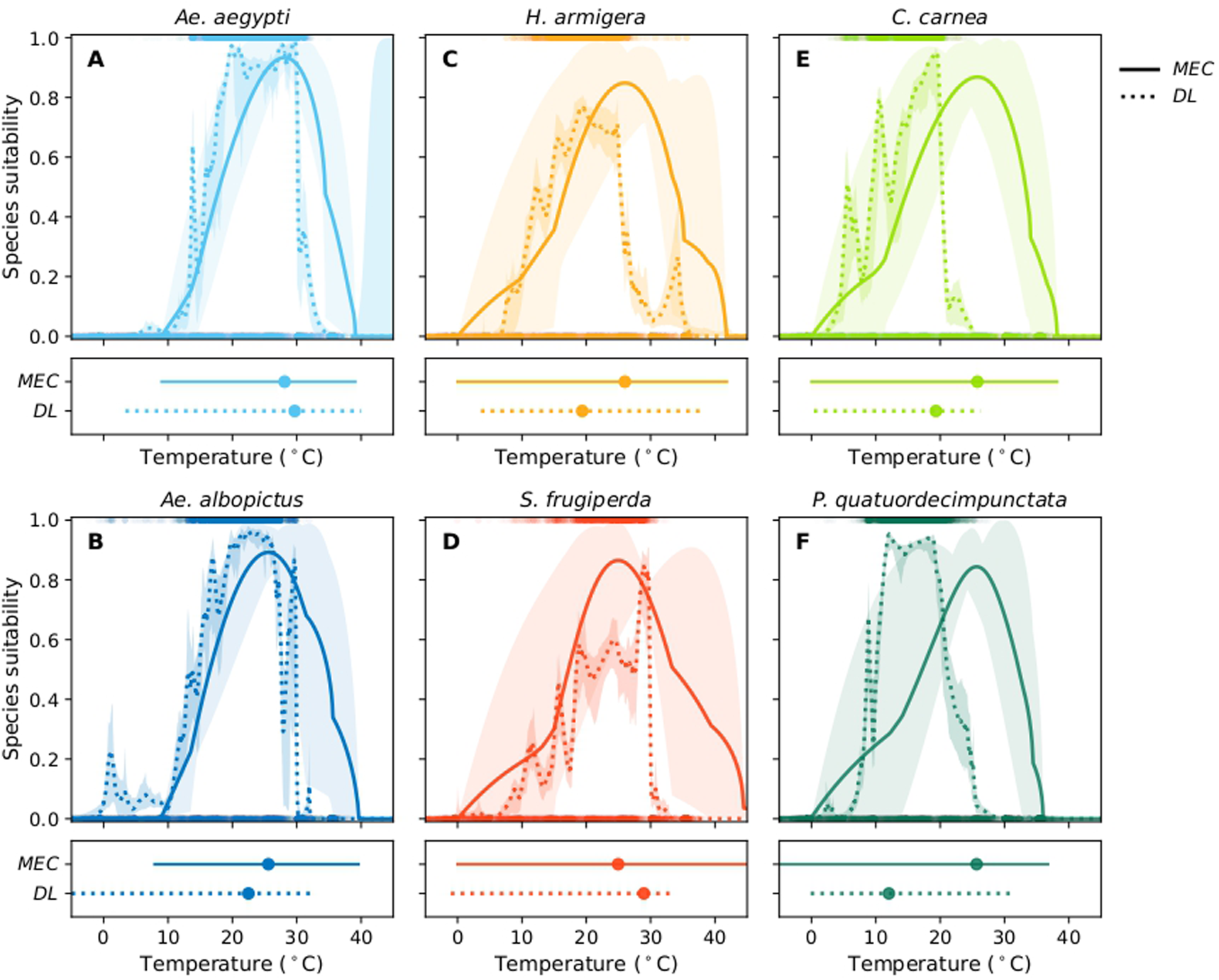
Species suitability along the thermal gradient, as estimated by mechanistic (solid line) and deep-learning (stratified background points, dotted line) approaches. (A) yellow fever mosquito (*Ae. aegypti*, in light blue), (B) asian tiger mosquito (*Ae. albopictus*, in blue), (C) cotton bollworm (*H. armigera*, in orange), (D) fall armyworm (*S. frugiperda*, in red), (E) common green lacewing (*C. carnea*, in light green), and (F) fourteen-spotted lady beetle (*P. quatuordecimpunctata*, in dark green). Below each graph, non-zero propbability areas (lines) and optimal temperatures (dots) are reported.

Deep learning-based suitability estimates, obtained with stratified background points, are less regular and suitability peaks follow the density distribution of species records across temperatures. They decrease drastically outside the observed thermal range (Figure 2). The latter is not always valid when deep learning-based suitability is trained using random background points (Figure A4). In this case, suitability might increase at extremely low (*Ae. albopictus* and *S.frugiperda*) or extremely high (*Ae. aegypti* and *H. armigera*) temperatures, overestimating the species’ suitable thermal range, whose extremes could not be identified. This behavior likely reflects unconstrained model extrapolation beyond the range of temperatures represented in the training data, resulting in suitability functions that are not ecologically robust. We therefore discard the use of random background points for further investigation of hybridization techniques. The *DL* approach with stratified background points presents thermal ranges and optima shifted towards lower temperatures, ranging from 12.0 to 29.7 *^◦^*C, specifically 29.7*^◦^*C or *Ae. aegypti*, 22.5*^◦^*C for *Ae. albopictus*, 19.4*^◦^*C for *H. armigera*, 28.9*^◦^*C for *S. frugiperda*, 19.3*^◦^*C for *C. carnea*, 12.0*^◦^*C for *P. quatuordecimpunctata*, respectively. For *Ae. aegypti* and *S. frugiperda*, although the *DL*-based thermal range is shifted toward lower temperatures, the estimated thermal optimum is slightly higher than the *MEC* one.

The highlighted differences in suitability estimates between *MEC* and *DL* approaches are further reflected in the derived distribution maps (Figure 3). *MEC* overestimates species suitability at high temperatures around the equator; these locations are not covered by species records (Figures 3 and A1). This is particularly evident for *C. carnea* and *P. quatuordecimpunctata*. Suitable areas identified by *DL*, and excluded by *MEC*, lie near the latitudinal extremes, which correspond to colder temperature conditions.

**Figure 3:**
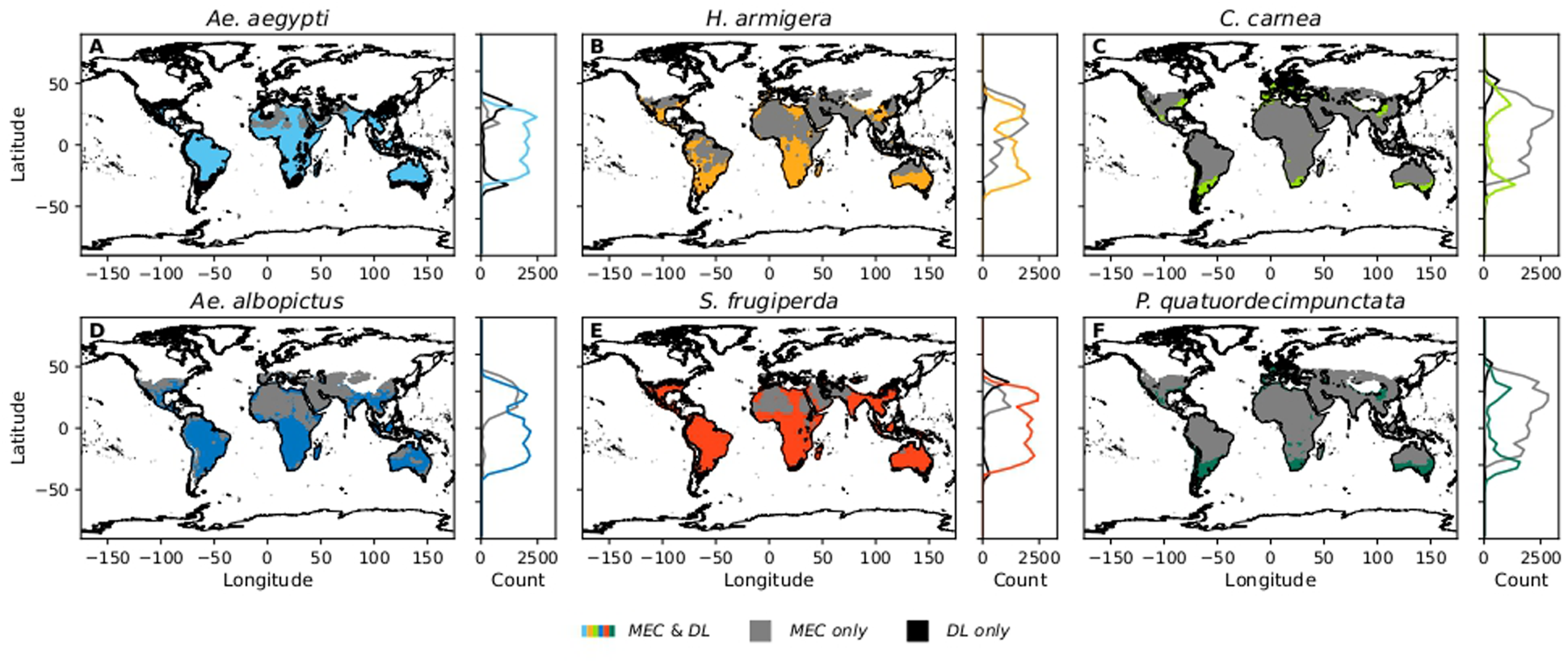
Distribution maps of (A) yellow fever mosquito (*Ae. aegypti*, in light blue), (B) asian tiger mosquito (*Ae. albopictus*, in blue), (C) cotton bollworm (*H. armigera*, in orange), (D) fall armyworm (*S. frugiperda*, in red), (E) common green lacewing (*C. carnea*, in light green), and (F) fourteen-spotted lady beetle (*P. quatuordecimpunctata*. Colored areas represent agreement between then *DL* and the *MEC* approaches, grey regions those identified as suitable only by *MEC* and the black ones only by *DL*. Beside each species suitability map, the latitudinal distribution of suitable areas is highlighted with the same color scheme.

When comparing the performance of the two approaches on the test set of species records, *DL* consistently and significantly outperforms *MEC* in terms of *AUC_ROC_* and *TSS* across all species (Figure 4). Performance gaps are smaller for the *Ae. aegypti* mosquito, for which *AUC_ROC_* is comparable across the two methods. The higher performance of *DL* is consistent with the greater similarity between the training and test tasks, which are both based on species occurrence, as well as with the ability of *DL* to capture more complex thermal functions, closer to natural conditions of thermal suitability. The *MEC* approach is less flexible and constrained by experimental temperature regimes, which explains the lower performance in predicting species’ realized distributions. Nonetheless, despite its lower predictive performance, *MEC* still provides valuable additional information by incorporating physiological responses to temperature that are not directly captured by *DL* models.

**Figure 4:**
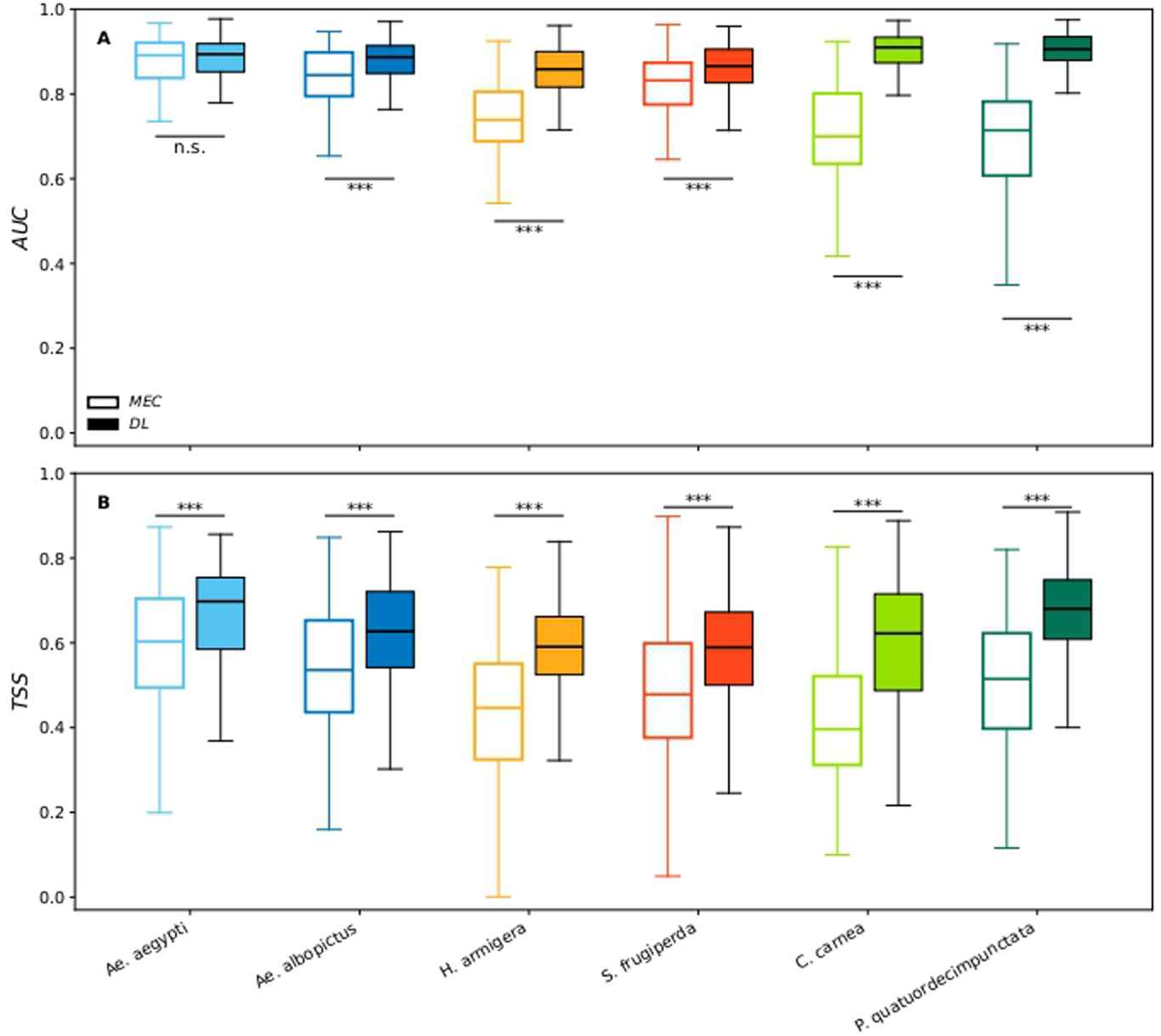
(A) Area Under the Receiving Operating Curve (*AUC_ROC_*) and (B) True Skill Statistics (*TSS*) performance scores on the species record test set (100 times resampling of spatial blocks) of the (*MEC*, empty boxplots) and deep learning-based (*DL*, filled boxplots) species distribution models for yellow fever mosquito (*Ae. aegypti*, in light blue), asian tiger mosquito (*Ae. albopictus*, in blue), cotton bollworm (*H. armigera*, in orange), fall armyworm (*S. frugiperda*, in red), common green lacewing (*C. carnea*, in light green), and fourteen-spotted lady beetle (*P. quatuordecimpunctata*, in dark green). Significance was assessed using a paired t-test comparing *MEC* and *DL* performance for each species. Asterisks indicate statistically significant differences (n.s.= *p ≥* 0.05*, ∗ ∗ ∗* = *p <* 0.001).

### Hybrid approaches and model robustness to data degradation

Exploiting full datasets of species records, *DL* seems to be better suited for modeling species distribution rather than experimentally based mechanistic approaches (Figure 5). However, the geographical distribution of species is rarely fully documented. When restricting available training data for the *DL* model to mimic potential data incompleteness while keeping the same test set for model evaluation (data degradation), we found a consistent drop in performance in the *DL* results (Figure 5). This decrease is more evident (Δ*AUC_ROC_*=[-0.45;-0.21]) when species records are degraded following the thermal gradient (Figure 5A, B), and is more abrupt when degradation occurs following the sampling density (Figure 5C). Conversely, when a random degradation is performed, *DL* shows more robustness, even up to the simulation retaining 20% of available data (Figure 5D). However, this data degradation procedure does not realistically represent existing biases regarding species distribution records.

**Figure 5:**
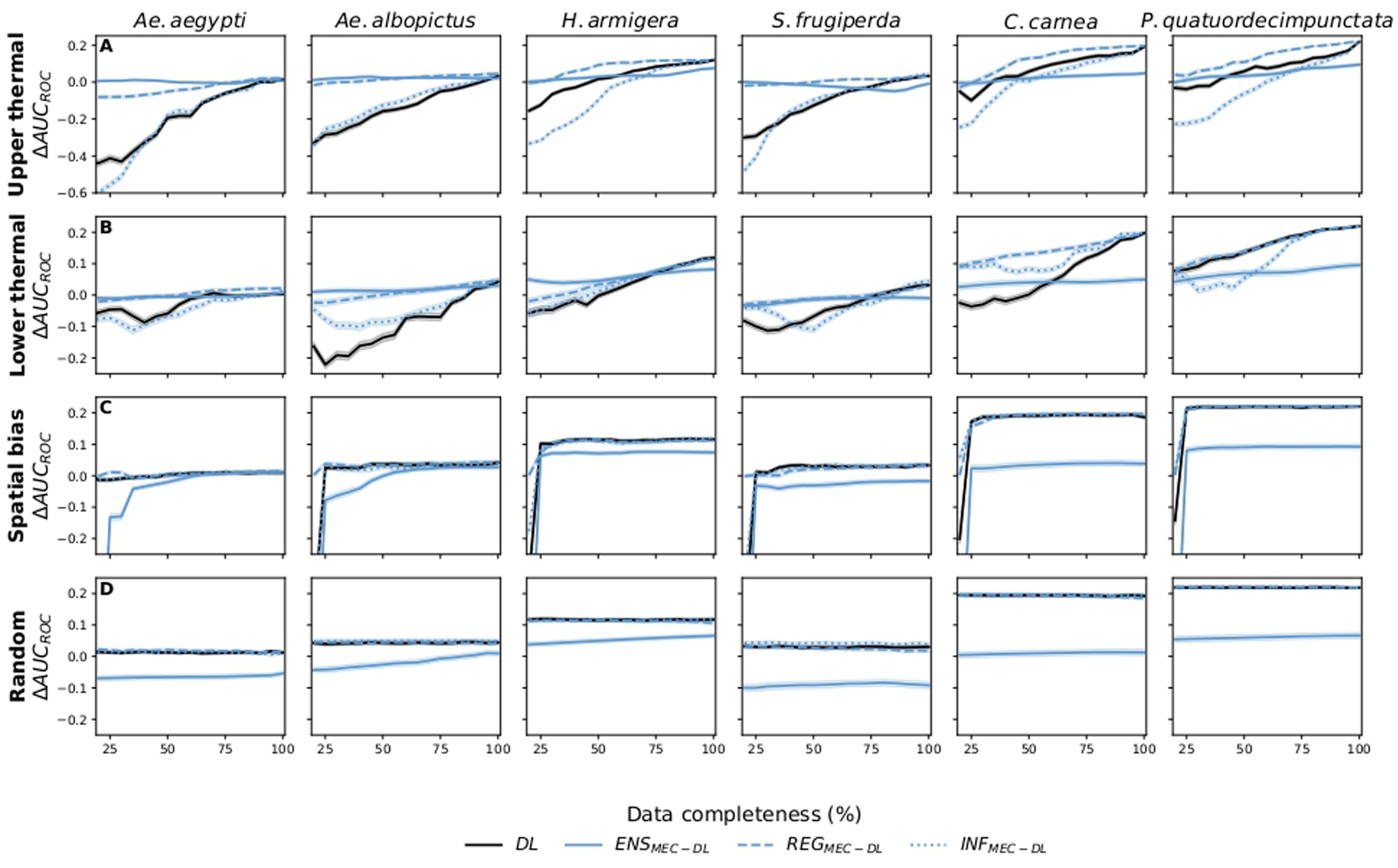
Species-specific differences in model performance (Δ*AUC_ROC_*) between deep learning-based suitability *DL* (in black) and hybrid approaches *MEC −DL* (in blue), compared to the mechanistic model *MEC*, whose performance is not affected by the degradation of the training set. Ensembling *ENS_MEC−DL_* (solid line), mechanistic regularization *REG_MEC−DL_* (dashed line), and mechanistically informed deep-learning *INF_MEC−DL_* (dotted line) are reported. The four rows show results for (A) upper thermal gradient, (B) lower thermal gradient, (C) spatial bias, and (D) random data degradation approaches. Shaded areas represent the standard error calculated from 100 runs of data resampling.

Hybrid approaches, in particular the model ensembling (*ENS_MEC−DL_*), as well as mechanistic regularization (*REG_MEC−DL_*) variants, were found to improve model robustness in case of data degradation. The drop in model performance is consistently less pronounced than for *DL* alone across all insect species and data degradation methods (Figure 5). Instead, the mechanistically informed variants *INF_MEC−DL_* seem to suffer from the same drawbacks of *DL*, with a persistent reduction in performance with degraded data along the thermal gradient. This result shows that it is not enough to pass the mechanistic suitability as an extra input to the *DL* model if the latter is not adequately considered in training. The hybridization approach that achieves the highest performance depends on the target insect species and the degree of data degradation applied. Generally, for data that are almost complete, mechanistic regularization appears to be the most effective approach. Conversely, for highly degraded data, mode ensembling tends to show a higher performance.

We selected 75% of data completeness according to the upper thermal degradation and explicitly compared the performance and thermal suitability of *DL* and mechanistic regularization *REG_MEC−DL_* methods (Figure 6). Consistently for all the analyzed species, the regularizing action of the mechanistic information results in higher model performance and more robust thermal suitability, whose extreme estimation is not constrained only by data availability in a certain climatic range. Importantly, even when the mechanistic model alone shows a limited ability to represent species records, it still brings complementary information that improves model adherence to species data (Figure 6C, E, F). Thus, hybridization frameworks, particularly model ensembling and mechanistic regularization, can improve predictive performance by integrating complementary knowledge from diverse sources, while making environmental function estimates less sensitive to sampling gaps and biases.

**Figure 6:**
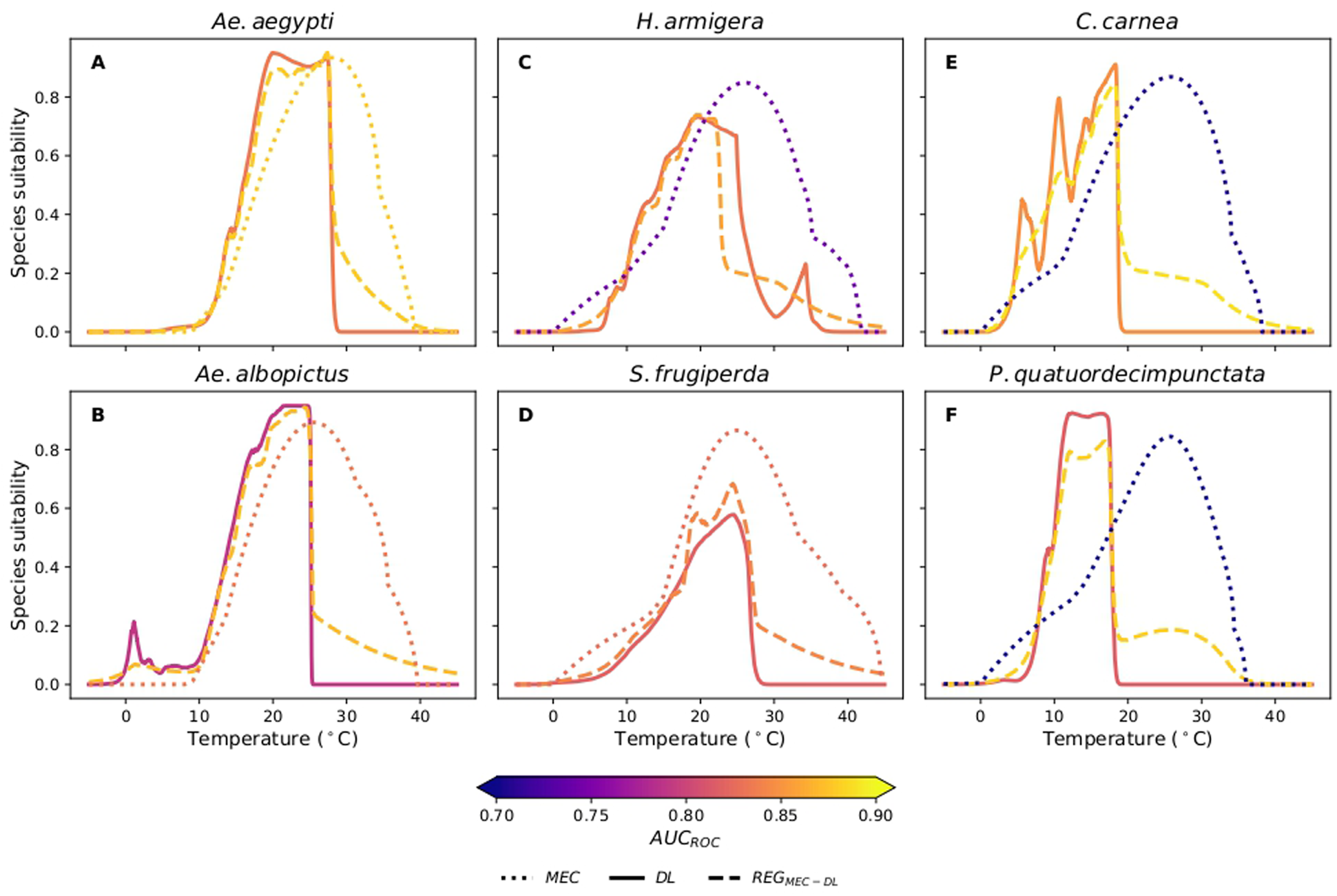
Species suitability along the thermal gradient estimated from 75% of data completeness according to the upper thermal degradation. Thermal suitability estimates are shown for the deep learning (*DL*, solid), mechanistic regularization (*REG_MEC−DL_*, dashed), and the original mechanistic (*MEC*, dotted) models. (A) yellow fever mosquito (*Ae. aegypti* ), (B) asian tiger mosquito (*Ae. albopictus*), (C) cotton bollworm (*H. armigera*), (D) fall armyworm (*S. frugiperda*), (E) common green lacewing (*C. carnea*), and (F) fourteen-spotted lady beetle (*P. quatuordecimpunctata*). The color of the lines corresponds to the *AUC_ROC_* performance of the respective model, as shown in the color bar.

## Discussion

Identifying insect species’ environmental responses is challenging, as climatic conditions affect the life stages of specimen in distinct ways, and because observations are often incomplete and potentially biased. In this study, we compared two complementary approaches: a process-based framework, grounded in experimental observations, and a correlative framework, based on species occurrences and realized thermal conditions. We evaluated three strategies for integrating deep learning with mechanistic environmental characterization: model ensembling, mechanistic regularization, and mechanistically informed neural networks. Model ensembling and mechanistic regularization improved robustness to data degradation, predictive performance, and the realism of environmental-response data-driven estimates under biased data. In contrast, näıve and mechanistically informed *DL* suffered from data degradation, with a strongly declining performance when data are degraded following the thermal gradient. These patterns were consistently observed for the analyzed disease insect vectors, agricultural pests, and biological control agents, despite their distinct thermal suitability and distributions.

Comparing species thermal suitability estimated from the process-based model with the correlative method, we found a consistent shift towards higher temperatures of the suitable ranges and thermal optima. This finding is potentially justified by the observed underfilling of the warmest part of the species’ niche for terrestrial species [75]. Thus, even if a location is thermally tolerable and could potentially be occupied, it gets avoided due to other abiotic and biotic constrains in the realized species distribution. The overestimation of lower thermal limit by the mechanistic suitability might also stem from thermal gaps in laboratory observations; additional experiments may provide more detailed information at temperatures below 10*^◦^*C (Figure A3). For instance, accurately identifying the lower thermal thresholds is particularly important for examining the potential for pole-ward expansion of a species’ niche under climate change [76]. Despite these limitations, the mechanistic approach does bring useful knowledge when hybridized with a correlative framework. This has been shown for the task of predicting gross primary productivity, *i.e.*, forest carbon assimilation by photosynthesis, using process-informed neural networks [25], or by integrating physiological trait species knowledge [77, 26]. In agreement with our findings, integrated mechanistic and correlative methods seem particularly useful in data-sparse regimes with high-transfer tasks [25], especially when data-derived knowledge is incomplete.

The value of species distribution models extends beyond establishing correlations between species records and environmental conditions. Through model-driven sampling design, they could potentially identify where additional sampling would be most informative, helping to allocate field efforts more efficiently and supporting more effective monitoring and ecological decision-making [78, 79]. This is particularly important because, even if species-specific process-based knowledge is considered universal in this study, ecological rules and patterns can vary across locations and change over time due to local adaptation and evolution. Therefore, integrating *DL* with adaptive sampling approaches can help to account for spatiotemporal variability, potentially suggesting where to sample insect colonies to further establish and refine trait-based lab experiments.

This study remains a proof of concept. Despite the demonstrated benefits of hybrid modeling, future research should focus on integrating additional environmental drivers, in addition to temperature, that highly impact insect ecology and distribution [15, 80]. For process-based approaches, this translates into extending the experimental design to incorporate additional abiotic conditions, which can substantially increase experimental time and costs. In contrast, integrating additional environmental drivers is relatively straightforward with correlative approaches and especially *DL*, as these models are inherently well-suited for handling multi-input and heterogeneous data sources describing species habitat [19, 21]. Following this path would require establishing a proper and independent test set for unbiased performance comparison across the multiple modeling approaches. Here, we used spatial block splits across species occurrence records from GBIF to examine model performance. This unavoidably skews the model comparison in favor of the correlative approach, bringing the test set closer to the task for which the *DL* model has been trained on. Consistent efforts have already been made in this direction by explicitly using different sources and protocols in constructing the training and test datasets, *e.g.*, leveraging the abundance of presence-only records for model calibration and independent, systematically collected presence–absence data from field surveys for model evaluation [81, 82, 83]. However, such structured presence–absence datasets remain geographically limited and are more available for plants [84] than animals, for which reliable absence data are particularly difficult to obtain.

In conclusion, addressing the knowledge gap regarding species environmental responses requires both bringing together complementary sources of ecological knowledge, and bridging correlative and process-based approaches in a combined modeling framework. While correlative models leverage the large volume of species occurrence data to infer species–environment relationships at broad spatial scales, process-based approaches provide mechanistic insights into the species physiological responses and thermal constraints. The rapid advances in artificial intelligence techniques have provided a lot of momentum towards the implementation of AI approaches in addressing ecological challenges [16]. However, studies like ours show that it is crucial to integrate these powerful methods with pre-existing knowledge of species ecology. In this context, a unified framework can facilitate the integration of knowledge traditionally developed across different disciplines and experts, enabling synergies unlocking new opportunities to advance our understanding of biodiversity.

## Data and Code Availability Statement

The data that support the findings of this study are openly available [64, 50, 51, 53, 52, 54, 54]. The data and code that support the findings of this study are openly available in a Figshare repository at https://figshare.com/s/2a4e46a55013f161b594.

