## Supplementary material for "Synergistic mechanistic and deep learning approaches for modeling insect thermal responses and geographic distributions"

### Appendix

Table A1: Literature references of the thermal dependence experimental observations of yellow fever mosquito (*Ae. aegypti*), asian tiger mosquito (*Ae. albopictus*), cotton bollworm (*H. armigera*), fall armyworm (*S. frugiperda*), common green lacewing (*C. carnea*), and fourteen-spotted lady beetle (*P. quatuordecimpunctata*) with respect to ecological traits (i.e., immature survival, fecundity, development, and adult lifespan). When available, the type of thermal performance curve is reported; otherwise, we trained on laboratory observations the two most common thermal functions (quadratic curve for adult survival and Brière function for the remaining traits).

| Species | Fecundity ($eggs/(female\ day)$ or $eggs/(female\ cycle)$) | Immature survival (%) | Development ($1/day$) | Adult lifespan ($day$) |
|---|---|---|---|---|
| *Ae. aegypti* | Brière [1] | Quadratic [1] | Brière [1] | Quadratic [1] |
| *Ae. albopictus* | Brière [1] | Quadratic [1] | Brière [1] | Quadratic [1] |
| *H. armigera* | Brière* [2] | Quadratic* [2] | Logan [2] | Quadratic* [2] |
| *S. frugiperda* | Brière* [3] | Extreme-value [3] | Brière [4] | Quadratic* [3] |
| *C. carnea* | Brière* [5] | Quadratic* [5] | Brière [6] | Quadratic* [5] |
| *P. quatuordecimpunctata* | Extreme-value [7] | Brière* [7] | Brière [7] | Quadratic* [7] |

*calibrated in this study from experimental observations

Table A2: GBIF species occurrence records after each filtering step: *i*) precise spatial coordinates (uncertainty < 1 $km$), *ii*) complete temporal information (month, year), and *iii*) records within activity season, both excluding occurrences during species-specific dormancy period.

| | *Ae. aegypti* | *Ae. albopictus* | *H. armigera* | *S. frugiperda* | *C. carnea* | *P. quatuordecimpunctata* |
|---|---|---|---|---|---|---|
| Raw occurrence | 27,188 | 72,736 | 20,521 | 2,947 | 12,981 | 39,926 |
| Precise spatial coordinates | 27,126 | 71,424 | 7,378 | 2,676 | 9,813 | 20,990 |
| Complete temporal information | 27,126 | 49,286 | 7,378 | 2,673 | 9,813 | 20,990 |
| Within activity season | 27,126 | 22,959 | 4,898 | 2,673 | 7,735 | 16,636 |

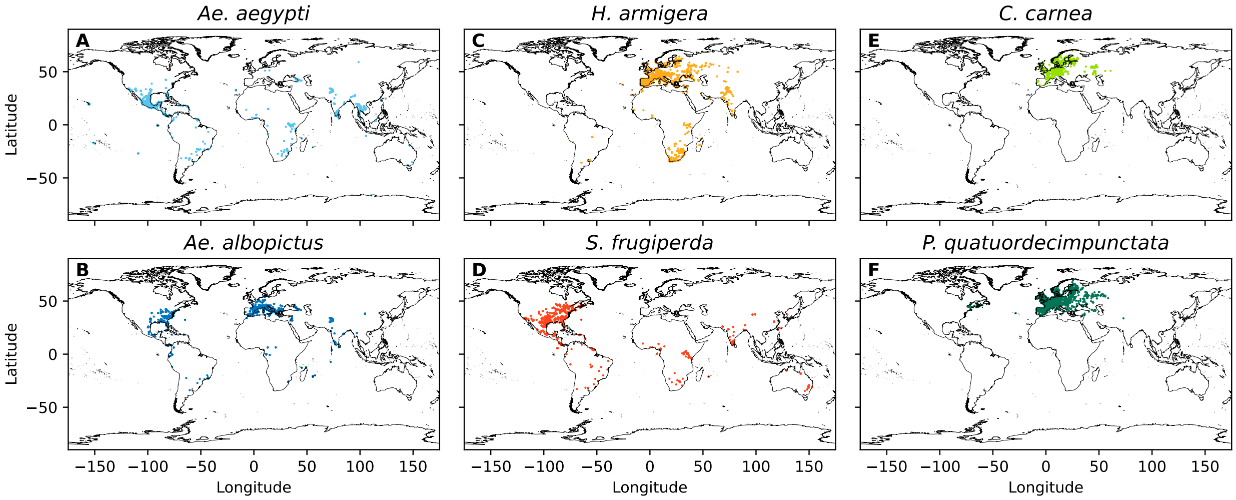


Figure A1: Insect occurrence records obtained from GBIF.

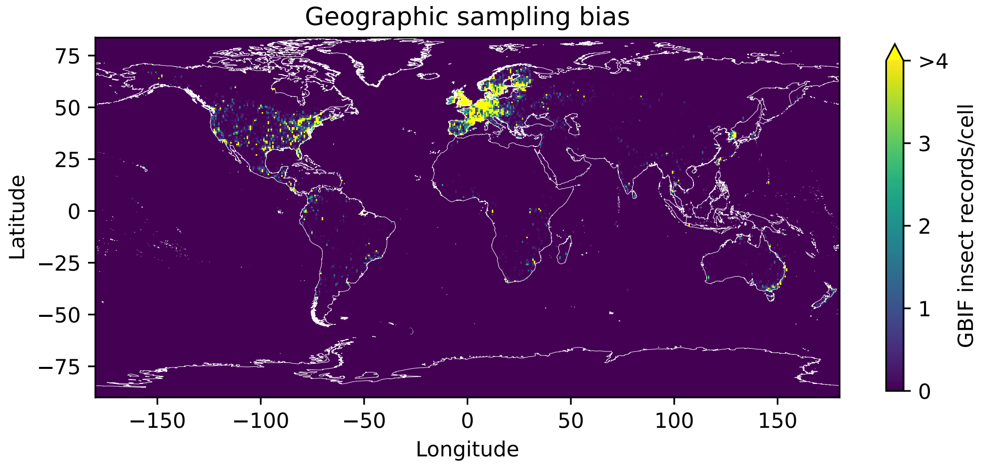


Figure A2: Global spatial density of insect occurrence records obtained from GBIF [8] for 1x1 $km^2$ spatial cells.

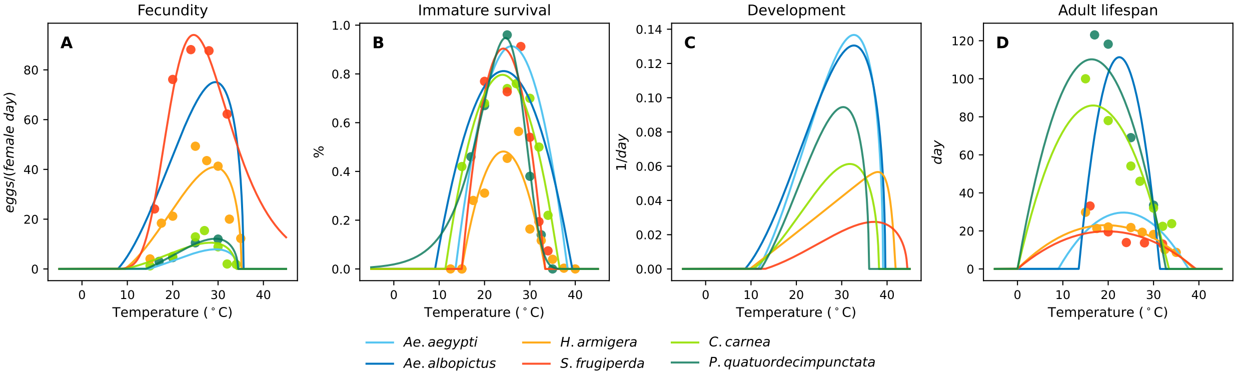


Figure A3: Temperature-dependent thermal performance curves for the ecological traits that define the mechanistic suitability model: (A) fecundity ($eggs/(female\ day)$ or $eggs/(female\ cycle)$), (B) immature survival (%), (C) development rate ($1/day$), and (D) adult lifespan ($day$). Lines represent the fitted thermal response functions and circles the experimental observations for *Ae. aegypti* (in light blue), *Ae. albopictus* (in blue), *H. armigera* (in orange), *S. frugiperda* (in red), *C. carnea* (in light green), and *P. quatuordecimpunctata* (in dark green). For details regarding selected thermal functions and literature references, see Table A1.

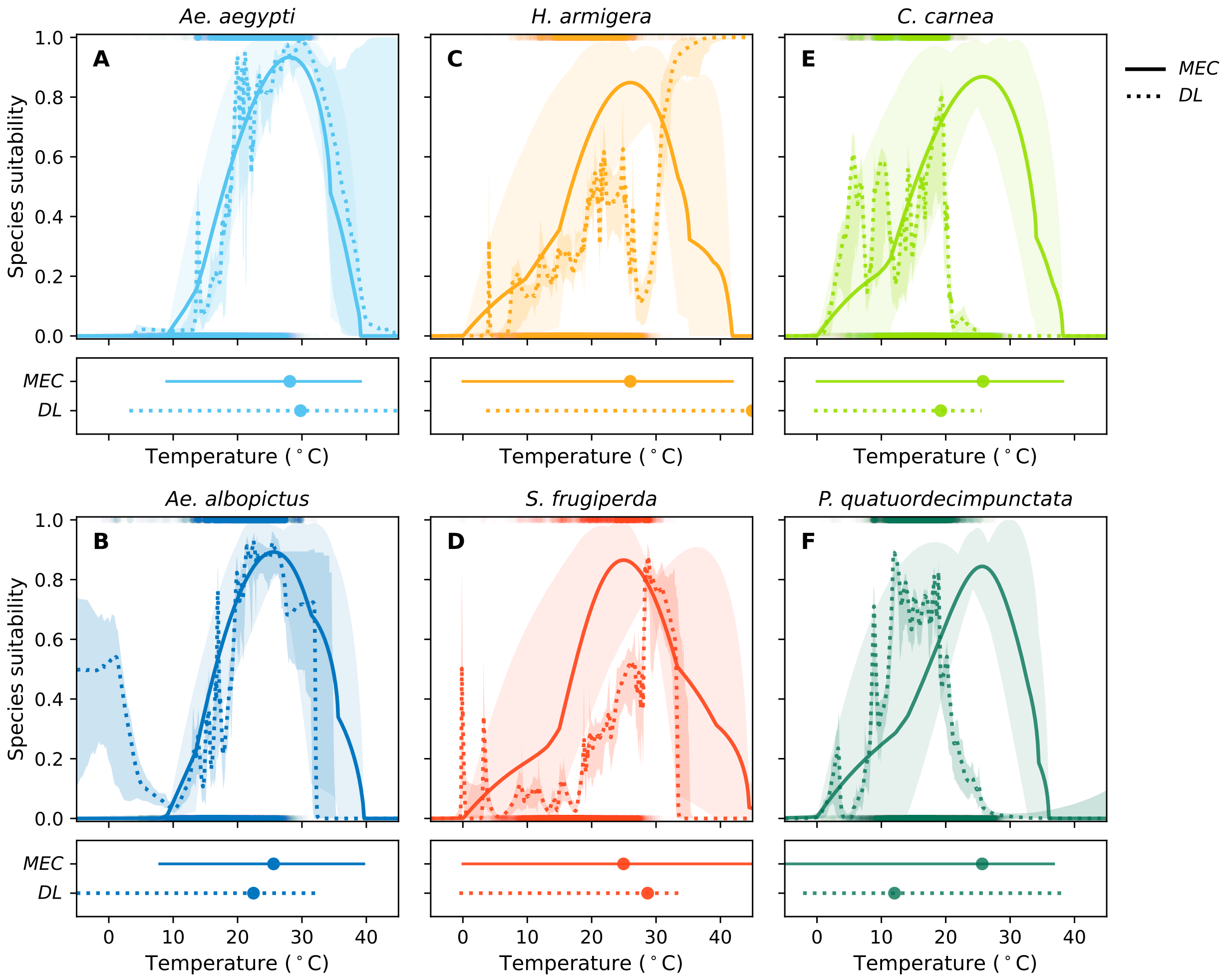


Figure A4: Species suitability along the thermal gradient estimated by the mechanistic ($MEC$, solid line) and deep-learning ($DL$ with random background points, dotted line) approaches for (A) yellow fever mosquito (*Ae. aegypti*, in light blue), (B) asian tiger mosquito (*Ae. albopictus*, in blue), (C) cotton bollworm (*H. armigera*, in orange), (D) fall armyworm (*S. frugiperda*, in red), (E) common green lacewing (*C. carnea*, in light green), and (F) fourteen-spotted lady beetle (*P. quatuordecimpunctata*, in dark green). Below each graph, non-zero propbability areas (lines) and optimal temperatures (dots) are reported.

Table A3: Sensitivity analysis of the mechanistic regularization weight $\lambda$. Model performance is reported as mean $\pm$ standard error (%) of the Area Under the Receiver Operating Characteristic Curve ($AUC_{ROC}$) for each species, for $\lambda \in \{0.1, 0.3, 0.5, 0.7, 0.9\}$. The highest mean in each column is highlighted in bold.

| Weight $\lambda$ | *Ae. aegypti* | *Ae. albopictus* | *H. armigera* | *S. frugiperda* | *C. carnea* | *P. quatuordecimpunctata* |
|---|---|---|---|---|---|---|
| $\lambda = 0.1$ | $89.61 \pm 2.02$ | $\mathbf{87.87 \pm 2.94}$ | $\mathbf{85.87 \pm 2.46}$ | $85.85 \pm 2.63$ | $\mathbf{90.05 \pm 2.40}$ | $90.64 \pm 1.61$ |
| $\lambda = 0.3$ | $89.77 \pm 2.09$ | $87.72 \pm 3.01$ | $85.61 \pm 2.56$ | $\mathbf{85.95 \pm 2.58}$ | $90.02 \pm 2.32$ | $\mathbf{90.66 \pm 1.57}$ |
| $\lambda = 0.5$ | $89.70 \pm 2.26$ | $87.62 \pm 3.04$ | $85.32 \pm 2.59$ | $85.47 \pm 2.73$ | $89.74 \pm 2.24$ | $90.43 \pm 1.63$ |
| $\lambda = 0.7$ | $89.99 \pm 2.03$ | $87.66 \pm 3.02$ | $84.86 \pm 2.63$ | $85.24 \pm 2.64$ | $89.32 \pm 2.18$ | $90.16 \pm 1.68$ |
| $\lambda = 0.9$ | $\mathbf{90.00 \pm 2.09}$ | $87.23 \pm 3.12$ | $83.91 \pm 2.83$ | $85.08 \pm 2.70$ | $88.72 \pm 2.22$ | $89.72 \pm 1.79$ |
